# Decoding the Amplitude and Slope of Continuous Signals into Spikes with a Spiking Point Neuron Model

**DOI:** 10.1101/2024.05.20.594931

**Authors:** Rebecca Miko, Marcus M Scheunemann, Volker Steuber, Michael Schmuker

## Abstract

Scalable and efficient neural models with biological plausibility are crucial for real-time applications in neuromorphic hardware and robotics. While biophysically detailed compartmental models are valuable for their accuracy, their complexity and computational cost limit real-world application on currently available hardware. In this study, we harness the signal processing potential of the Izhikevich point neuron model to decode slope and amplitude, both important dynamical features of sensory signals. Results demonstrate that our slope-detector effectively signals up-stokes of naturalistic input signals across a wide range of frequencies. It exhibits bidirectional slope detection, with burst duration encoding slope magnitude in a graded manner. We compare to a biophysically detailed two-compartment pyramidal neuron model, showing that our bursting slope-detector performs similarly. We then demonstrate that our slope-detector does not need to burst, improving efficiency and producing more precise, discrete output by signalling events with single spikes. This makes it well-suited for real-time robotics control systems or neuromorphic hardware applications, offering greater efficiency and enabling large-scale simulations using the same computational power.

## Introduction

In various research fields like gas-based navigation in robotics, computationally efficient models that balance biological realism and can efficiently process information is a requirement. Amplitude and slope are important dynamical features of sensory signals, encoding information about the scene [12, 14, 15]. The instantaneous dynamics of gas concentration in a turbulent plume are extremely complex [4, 12], and this rich temporal structure and spatial distribution of gas plumes demand rapid, low-latency responses to temporal cues in these signals [4, 13, 22].

Despite great strides in the development of neuromorphic hardware, such as the 2nd generation SpiNNaker [36] and Loihi [19], implementing complex neural models incurs higher computational costs [18, 24, 35]. We create amplitude and slope-detectors from the Izhikevich point neuron model [7], which inherently processes information with less computation in lower-dimensional space. The model maintains an effective trade-off between the biological plausibility of Hodgkin-Huxley-type dynamics and the computational efficiency of Integrate-and-Fire neurons, making it a popular choice for neuronal computations on neuromorphic hardware [26, 32, 33] and robots [21, 28, 34].

Izhikevich presented a two-dimensional system of ordinary differential equations (Eq. 1) with an auxiliary after-spike reset (Eq. 2). The parameters in the equations map to known biological features of neuronal spiking processes.

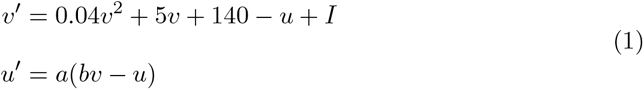

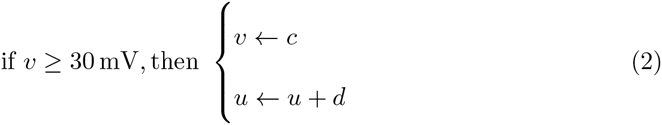

The dimensionless variables, membrane potential (*v*) and recovery variable (*u*), represent the core elements of the model. The parameters (*a, b, c, d*) control the recovery rate of *u*, sensitivity of *u*, after-spike reset of *v*, and after-spike reset of *u*, respectively. The model’s mathematical simplicity, with only one non-linear term (*v*^2^), contributes to its computational efficiency.

Though the model lacks specific biophysical mechanisms and spatial complexity, its adaptation term allows it to replicate various neural behaviours, including intrinsic bursting and fast-spiking. Adaptation also allows the neuron to operate under a certain degree of independence from the absolute magnitude of the input signal, a prerequisite for responding to changes in the signal’s derivative, i.e., detecting slopes, rather than merely reacting to changes in input strength. Other widely used models, such as the Leaky Integrate-and-Fire (LIF) neuron model [6], also offer computational efficiency and can be effectively integrated onto neuromorphic hardware [20, 25, 31]. However, the absence of an adaptation term makes slope-detection unattainable [10, 11].

In this study, we discover amplitude and slope-detectors, through simulations with sinusoidal and naturalistic input currents. We test our bursting slope-detector’s ability across a wide range of frequencies and demonstrate its bidirectional slope detection. We then show how its burst duration encodes slope magnitude in a graded manner. These results provide valuable insights into how these neurons can be utilised for real-time processing in dynamic, complex environments and tasks like gas plume tracking.

We later compare our bursting slope-detector to a two-compartmental biophysical neuron model, showing how the two models perform similarly, with one key difference. The model by [5] includes a dendrite compartment with persistent sodium and slow potassium currents responsible for bursting behaviour, while the somatic region contains Hodgkin-Huxley-type [1] currents that generate fast spikes. The authors showed how it spikes on the rising edges of naturalistic inputs signals. Our study shows that, unlike such complex models, regular spiking point neurons can effectively function as slope-detectors, making the bursting mechanism unnecessary for slope detection while still exhibiting adaptation in their behaviour. These neurons have less complex dynamics, making them ideal for applications which require computational efficiency.

## Results

We present results from our search of the Izhikevich parameter space, identifying optimal sets for amplitude and slope-detectors. We then analyse specific detectors. Finally, we compare our results to the biophysically detailed two-compartment pyramidal neuron model [5].

### Identifying slope and amplitude-detectors

We present our search results for optimal detectors in the Izhikevich parameter space. Three identical grids are shown, each covering a wide range of values for the parameters *a, c*, and *d* (see Eq. 1). The grids display the slope detection percentage, amplitude detection percentage, and bursting percentage for each parameter combination.

Fig. 1a presents the grid for slope detection. It shows that low values of parameter *a*, particularly in combination with higher values of *c* (such as *c* = −35), produced strong slope-detecting behaviour. Based on this observation, we selected the neuron with parameters {a:0.01, b:0.2, c:-35, d:5.0} as an example slope-detector (Fig. 1b).

**Fig 1.**
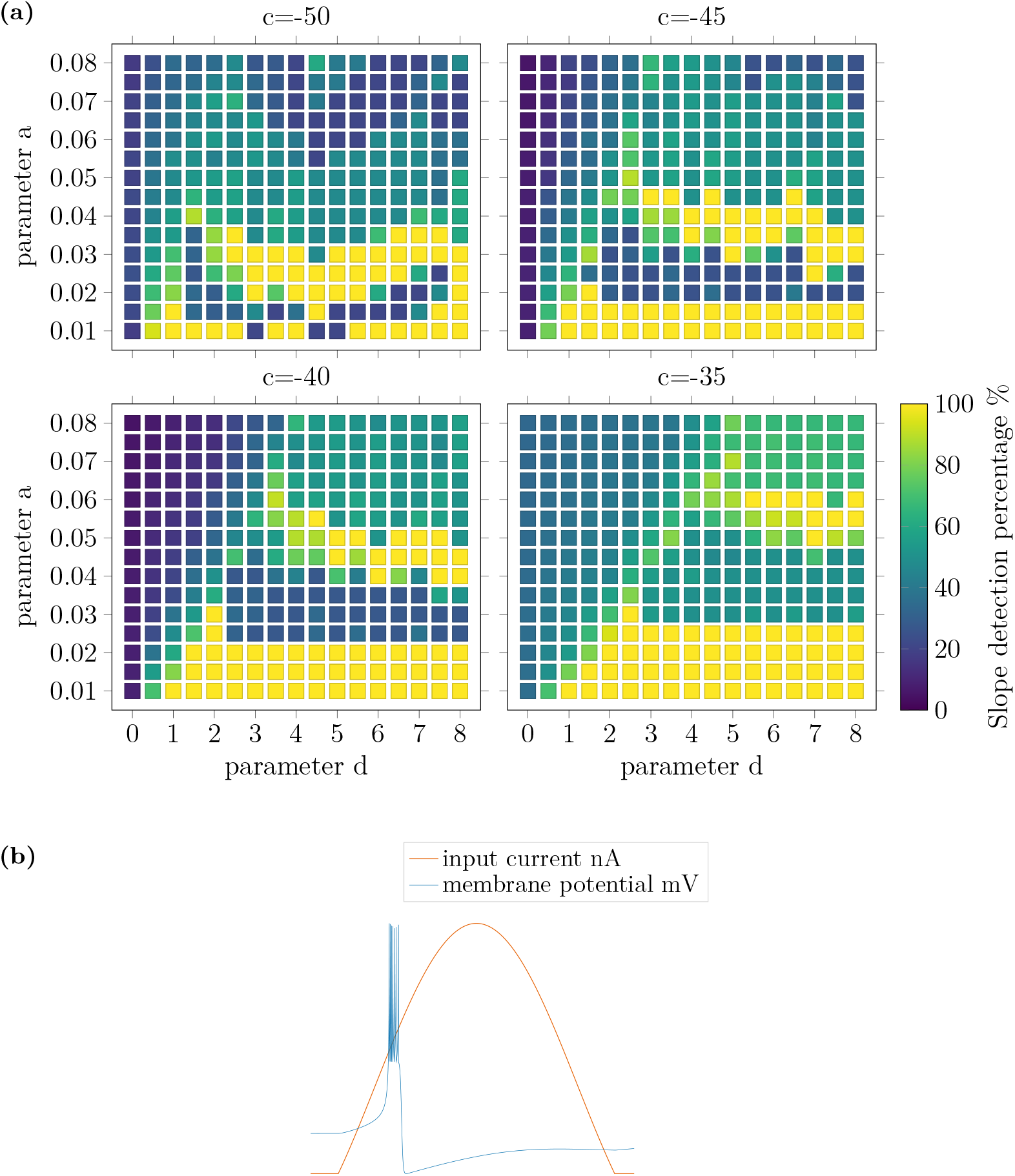
Results in **(a)** show parameter combinations that enable slope-detection; Izhikevich parameters *d* (x-axis), *a* (y-axis), and *c* (panels). Slope-detection capability is confined to specific regions in parameter space. Slope detection percentage highlights regions that generate strong slope-detectors (yellow). Depicted in **(b)** is the membrane potential of a slope-detector ({a:0.01, b:0.2, c:-35, d:5.0}) and a rectified segment of the 4 Hz sinusoidal input signal. A burst of spikes on the rising edge shows slope-detection.

Fig. 2a shows the grid for amplitude detection. It shows that lower values of *d* combined with higher *c* values (such as *c* = −35) produced strong amplitude-detecting behaviour. Higher *c* values also generate a larger area of slope-detecting neurons, suggesting a commonality with the membrane voltage reset.

**Fig 2.**
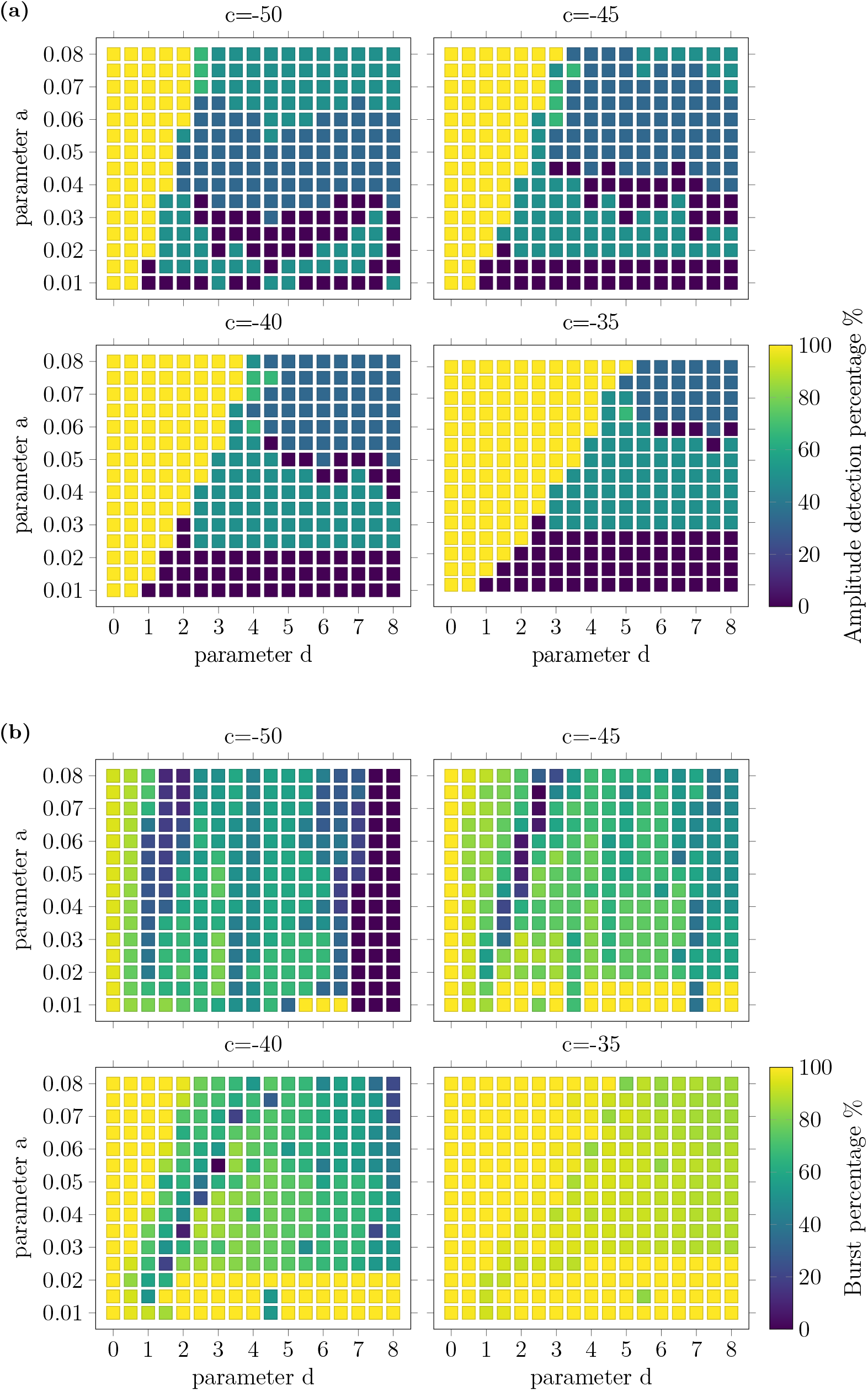
Results in **(a)** show parameter combinations that enable amplitude-detection; Izhikevich parameters *d* (x-axis), *a* (y-axis), and *c* (panels). Amplitude-detection capability is confined to specific regions in parameter space. Detection percentage highlights regions that generate strong detectors (yellow). Results in **(b)** show parameter combinations that enable bursting, defined by an inter-spike interval of 10 ms.

The comparison between the slope-detector grid (Fig.1a) and the amplitude-detector grid (Fig.2a) demonstrates cases where a set of parameters can produce both slope and amplitude-detecting neurons. For instance, the neuron with parameters {a:0.04, b:0.2, c:-35, d:5.0} exhibits a slope detection percentage and an amplitude detection percentage of 50 %, indicating that it spikes on both the rising edges and signal peaks.

Fig. 2b shows the grid for bursting behaviour. Higher *c* values generally lead to bursting, while the combination of parameters *a* and *d* also affects the bursting mechanisms. Furthermore, we observe no strong relationship between the bursting grid (Fig.2b) and the slope (Fig.1a) and amplitude (Fig. 2a) detector grids.

We then test the slope and amplitude-detectors by stimulating them with temporal input signals, created by filtered Gaussian white noise. The results (Fig. 3) show the consistent and accurate encoding of either slope or amplitude. Fig. 3a shows that the slope-detector bursts preferentially at rising edges of the input signal, while Fig. 3b shows that the amplitude-detector bursts preferentially at signal peaks.

**Fig 3.**
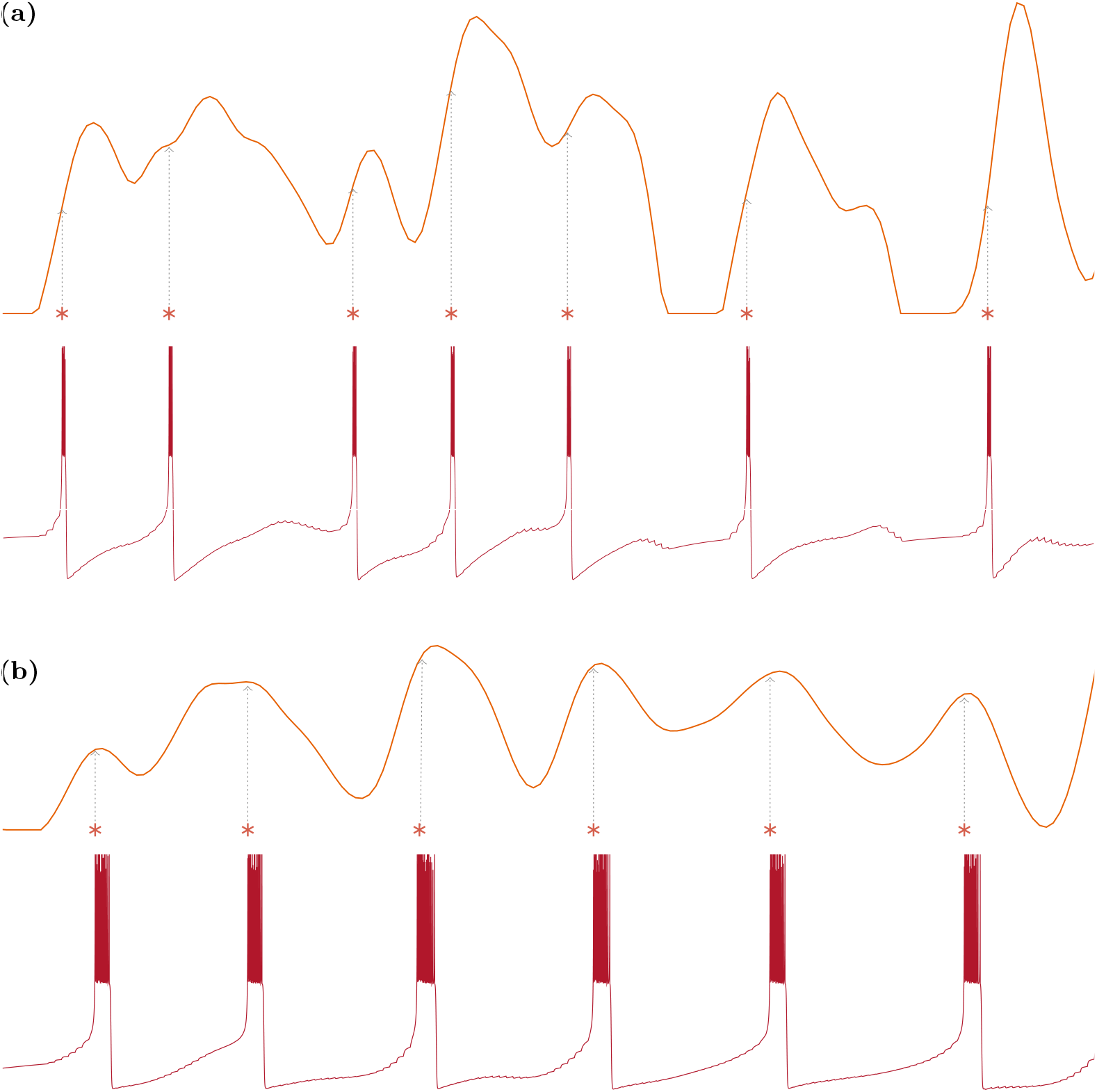
*Top trace*: filtered Gaussian white noise input nA (5 Hz; *µ* = .006; *σ* = .015). *Bottom trace*: membrane potential mV response. Results in **(a)** and **(b)** show the neurons with parameters {a:0.01,b:0.2,c:-35,d:5.0} and {a:0.05,b:0.2,c:-40,d:1.0}, respectively. *Asterisks* mark burst onsets (*grey dotted lines* for clarity). Bursts, defined by an inter-spike interval of 10 ms, occur at current peaks. No single spikes produced.

### Detection frequency range

We analyse the response of the bursting slope-detector to stimulus upstrokes at different frequencies using the Spike-triggered Average (STA) plot, as shown in Fig. 4. This shows the average stimulus that precedes the neuron’s spikes, revealing the features of the stimulus that are most likely to trigger neuronal firing.

**Fig 4.**
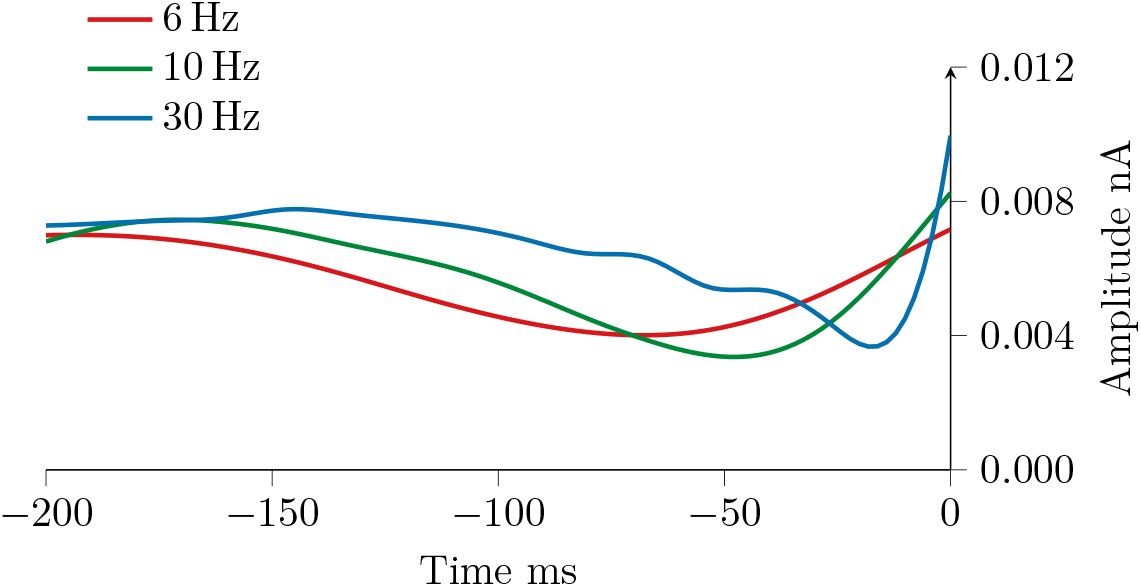
Results of the Spike-Triggered Average (STA) analysis, showing input signal features that trigger bursts. The bursting slope-detector neuron with parameters {a:0.01,b:0.2,c:-35,d:5.0} follows stimulus upstrokes over a wide range of frequencies.

The STA plot demonstrates that this detector is capable of responding to stimulus upstrokes across a wide range of frequencies.

### Bi-directional slope detection

The results reveal that bursts occur on the upstrokes of the signal (e neuron) and on the down strokes of the inverted signal (i neuron), as demonstrated in Fig. 5, showing that the slope-detector is capable of bi-directional slope detection.

**Fig 5.**
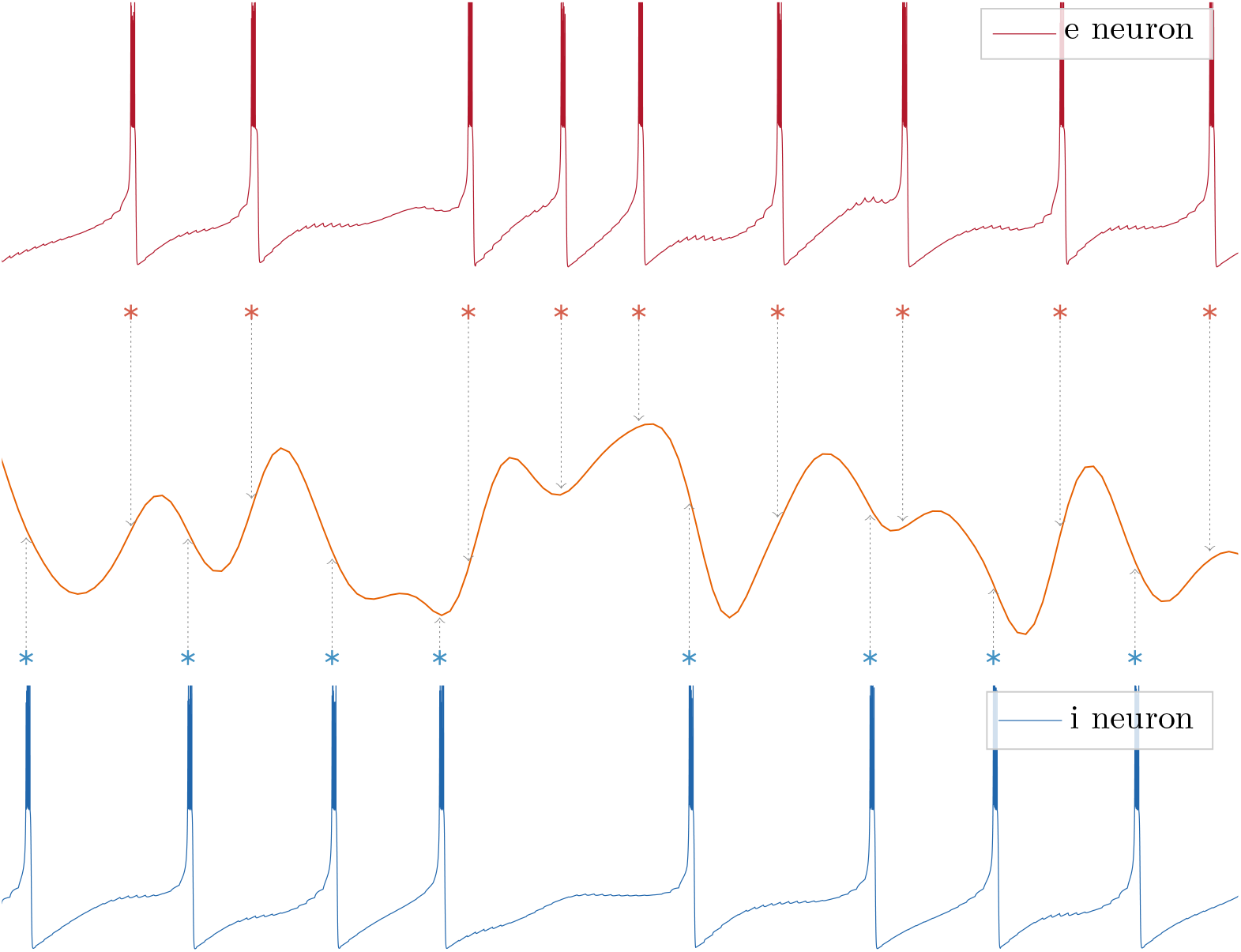
*Middle trace*: filtered Gaussian white noise input nA (5 Hz; *µ* = .008; *σ* = .015). *Top trace*: membrane potential mV response from e neuron. *Bottom trace*: membrane potential mV response from i neuron, where the *middle trace* is inverted. *Asterisks* mark burst onsets (*grey dotted lines* added for clarity).

### Burst duration encoding

Our studies show that the detector not only marks the occurrences of slopes in the signal but also encodes the magnitude of input slopes in a graded manner through the length of bursts, indicated by the number of spikes (Fig.6a, Fig.6b).

**Fig 6.**
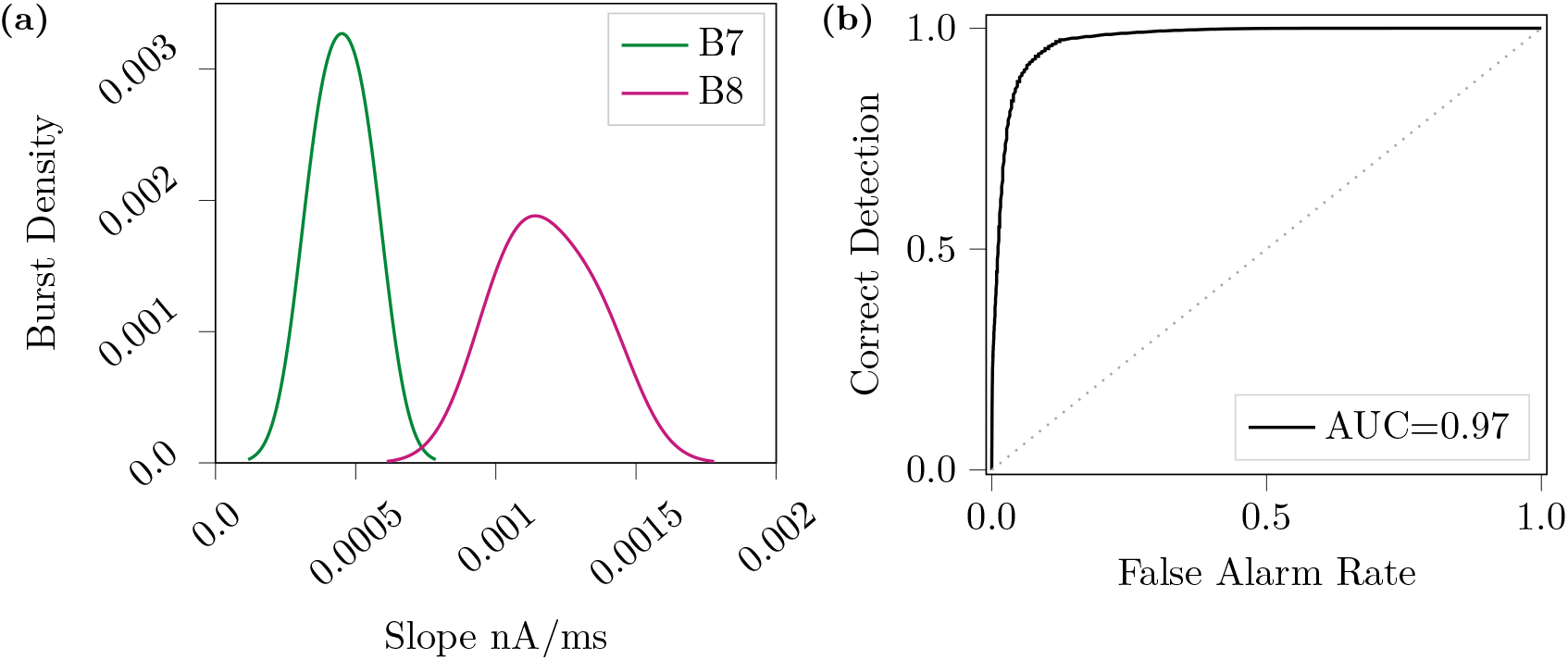
Distributions of input slopes for bursts durations in **(a)**, where the neuron with parameters {a:0.06,b:0.2,c:-35,d:5.5} produces 7- or 8-spike bursts. Receiver Operator Characteristics (ROC) curve in **(b)** shows the discriminability of 7- and 8-spike burst distributions. The Area Under the Curve (AUC) value is included for clarity.

We increase the variable *a* from 0.01 to 0.06, resulting in the neuron producing bursts of 7- or 8-spikes (*B7, B8* in Fig.6a). These burst durations correspond to different slope magnitudes with little overlap, as indicated by the Area Under the Curve (AUC) value of 0.97 for the Receiver Operating Characteristic (ROC) curve in Fig.6b.

### Bursting slope-detector comparison

We compare the bursting slope-detector neuron with the biophysical neuron model proposed by Kepecs et al. [5]. We generate similar input signals and analyse the output responses to evaluate the performance of the detector. The evaltuions in the section are qualitative.

Fig. 7 shows the membrane potential response of the bursting slope-detector neuron to a filtered Gaussian white noise input signal (5 Hz; *µ* = .008; *σ* = .015). The results show that the bursting slope-detector neuron behaves similarly to the biophysical model, sharing the ability to signal consecutive upstrokes without intervening down strokes. Although, the detector produces a cleaner and more efficient output, as there are no isolated spikes. Spikes are more selectively triggered by meaningful input features, responding specifically to upstrokes and avoiding redundant firing.

**Fig 7.**
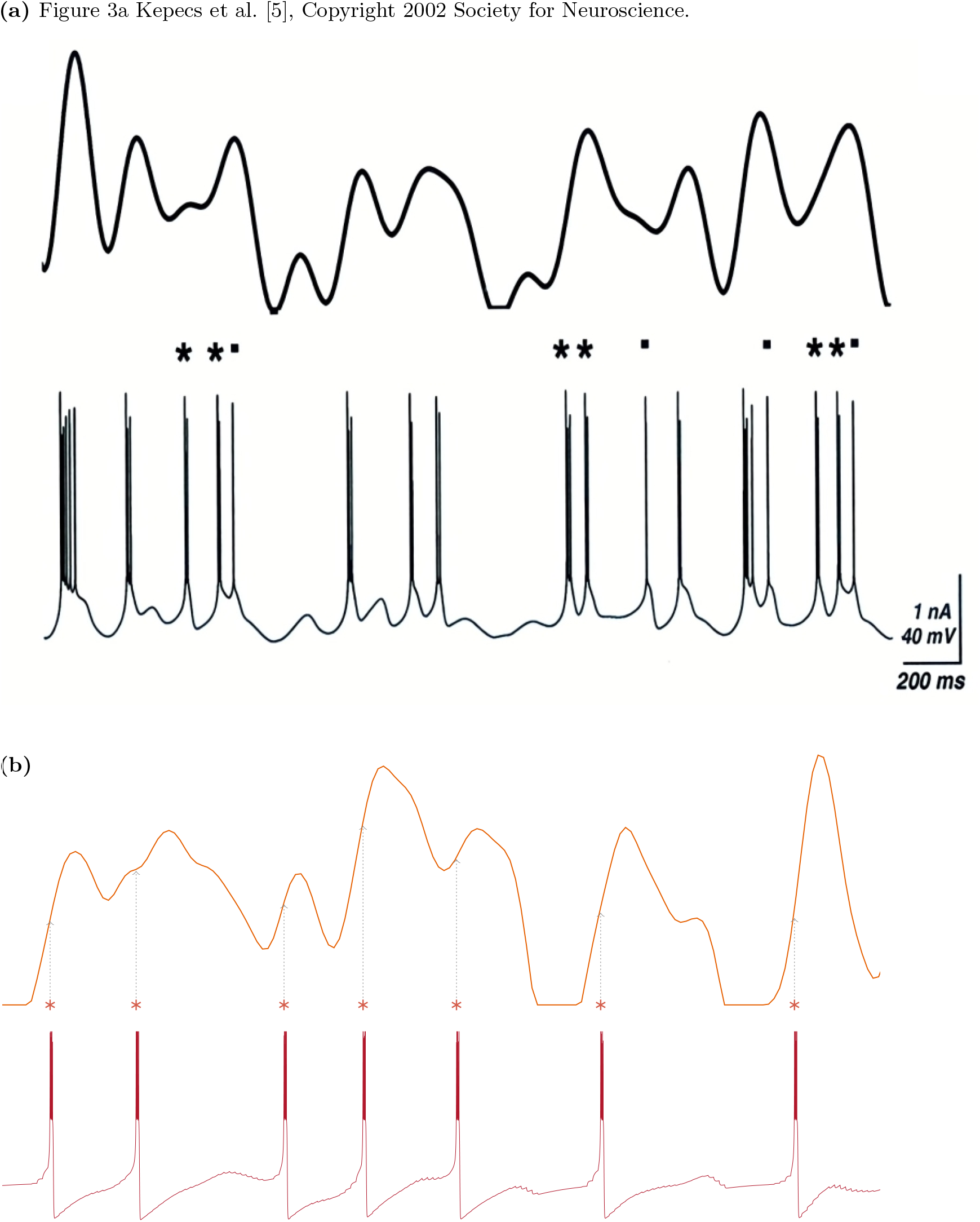
Relationship between input signal nA (*top trace*) and membrane potential mV response (*bottom trace*). *Asterisks* mark burst onsets. Bursts are defined by an inter-spike interval of 10 ms. *Dots* mark single spikes. The image in **(a)** is Fig. 3a in Kepecs et al. [5], Copyright 2002 Society for Neuroscience. Input (with 5 Hz; *µ* = 1.7; *σ* = 1.0) was applied to the dendrite and bursts were triggered on the positive slopes. For our study, **(b)** shows results from the Izhikevich bursting slope-detector with parameters {a:0.01,b:0.2,c:-35,d:5.0}. Filtered Gaussian white noise input signal nA (with 5 Hz; *µ* = .006; *σ* = .015) is injected. *Grey dotted lines* are added for clarity, showing bursts are triggered on the positive slopes. No single spikes are produced.

We found that while our bursting slope-detector is comparable to the biophysical neuron model, an Izhikevich slope-detector does not need the bursting mechanism to encode input slopes. The neuron with parameters {a:0.01,b:0.2,c:-50,d:8.0} spikes at the rising edges of a comparative signal (Fig. 8). The spike rate alone can indicate input slopes and the detector is not limited to sinusoidal input signals. This means that even without a complex bursting mechanism, but while still exhibiting adaptation in their behaviour, an Izhikevich point neuron can encode input slopes. This offers a computationally efficient and biologically plausible way to detect dynamic signal changes across multiple input types. This creates a digital, binary-like output where spikes indicate signal slopes and the absence of spikes reflects no relevant features are detected. A practical implication is that this clean signal makes it easier to integrate our neuron onto a digital system, such as a robot.

**Fig 8.**
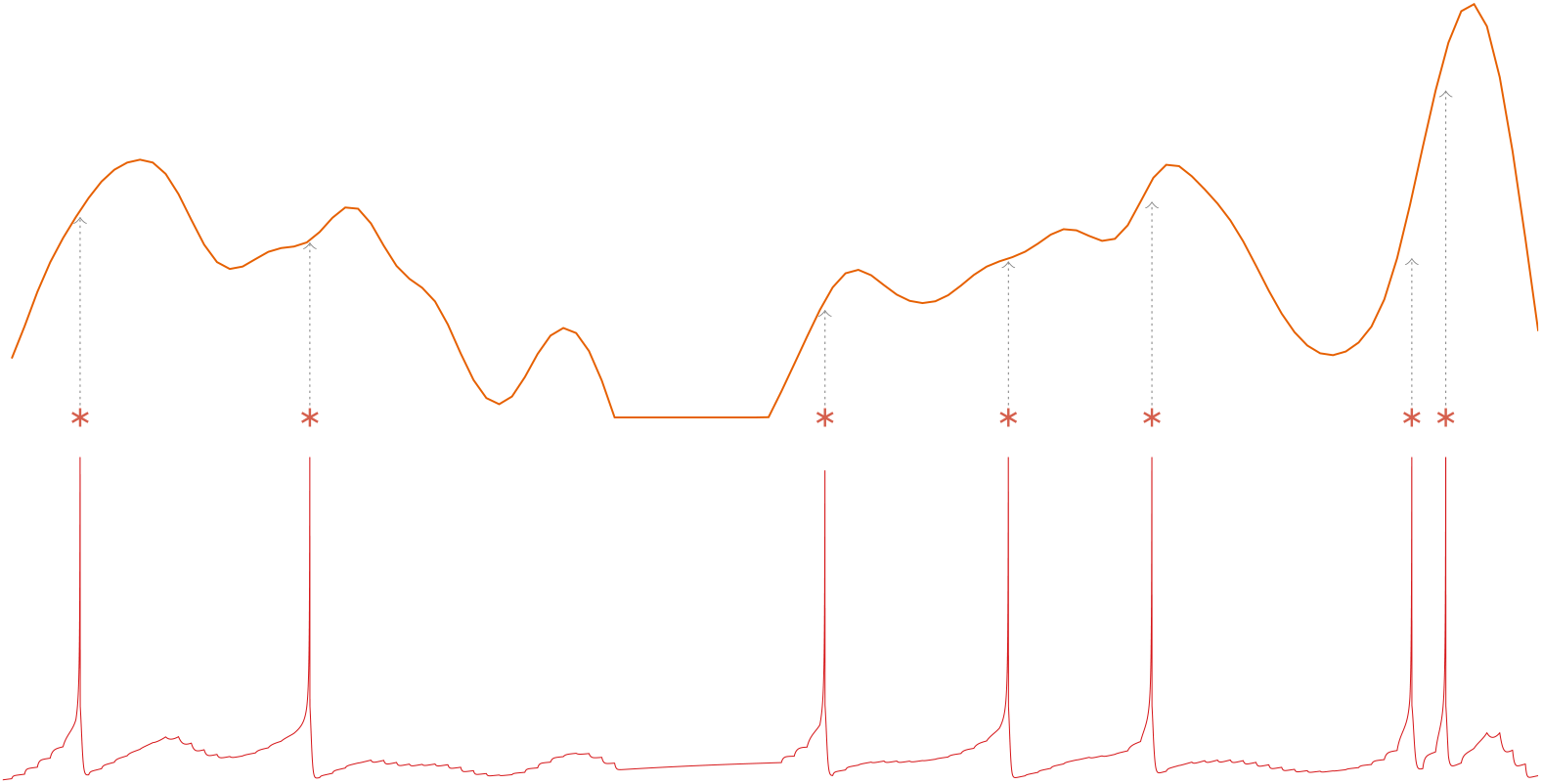
*Top trace*: filtered Gaussian white noise input nA (5 Hz; *µ* = .006; *σ* = .015). *Bottom trace*: membrane potential mV response. Slope-detector neuron with parameters {a:0.01,b:0.2,c:-50,d:8.0}. *Asterisks* mark spikes. *Grey dotted lines* are added for clarity, showing spikes are triggered on the positive slopes of the input signal.

We further compare the models qualitatively by looking back at our previous results for the slope-detector neuron. The Spike-triggered Average (STA) analyses can be compared to the STA from Kepecs et al. [5] to find that both neurons can encode the slopes of a comparable input signal within the same range of frequencies. The biophysical neuron model also demonstrated bidirectional slope detection.

Regarding burst duration encoding, both the slope-detector and biophysical neuron model encode the magnitude of input slopes in a graded manner. Both define a burst by an inter-spike interval of 10 ms, and show that burst duration increases with steeper input slopes. This graded response reflects the sensitivity of the neuron to the rate of input change rather than input amplitude.

The models differ here in the number of spikes in a burst, as the biophysical neuron has burst lengths of 2- to 5-spikes and the slope-detector has burst lengths of 7- to 8-spikes. The low reset potential (*c* = −35) and moderate recovery parameters (*a, b, d*) create conditions for prolonged bursting in response to increasing input slopes. This makes the slope-detector neuron more responsive to sustained depolarization, encoding slope magnitude with longer burst lengths. This difference highlights that the two-compartment biophysical model, with its four distinct burst lengths as opposed to two, can represent a broader and more nuanced range of slope magnitudes.

## Discussion

The main aim of this work was to determine whether a computationally efficient and biologically plausible point neuron model could decode key dynamical features of sensory input in a manner suitable for future real-time applications in robotics and neuromorphic hardware. Our study demonstrates how to create amplitude and slope-detecting neurons, using the Izhikevich point neuron model. We demonstrate that simple Izhikevich neurons can reliably signal the rate of change in naturalistic inputs, initially through bursting dynamics similar to those observed in a biophysically detailed two-compartment pyramidal neuron model [5]. We then demonstrate that even regular spiking point neurons can function effectively as slope-detectors, enabling discrete and efficient event signalling. This makes the bursting mechanism unnecessary for slope detection while still exhibiting adaptation in their behaviour.

This work lays the foundation for applying slope-detecting neurons to various research fields, particularly in the processing of sensory signals where amplitude and slope are key dynamical features that encode information about the scene [12, 14, 15]. The ability to detect both upstrokes and down strokes of the input signal enables potential applications in gas-based control research [4, 14, 16, 23, 30], where detecting the presence and absence of gas provides valuable information about the olfactory scene. The ability of the model to report the magnitude of input slopes in a graded manner through the length of bursts produced further supports an application in gas-based control studies, where the slope durations can inform the distance to an odour source [14]. These results enable the effective implementation of gas-based control in mobile robot applications, expanding the experimental capabilities for studying navigation in dynamic olfactory environments on embedded hardware.

We demonstrate that a regular spiking point neuron can function effectively as a slope-detector. While regular spiking neurons lack the bursting mechanism that allows for magnitude encoding, they offer key advantages over bursting neurons such as being more computational efficient. They create a discrete, binary representation of the signals where spikes indicate signal slopes and the absence of spikes reflects no relevant features are detected. Practically, this clean signal makes it easier to integrate our neuron onto a digital system, such as a robot. This is because all interpretation is already compacted, and the robot’s control system can directly act on events detected by our system. Regular spiking neurons have less complex dynamics, while still exhibiting adaptation in their behaviour, making them ideal for large-scale simulations and real-time applications [7]. Regular spiking neurons are more easily integrated into larger computational models [6].

### Impact and future work

The broader impact of this study extends beyond simple bio-mimicry, it presents a complementary approach in neuromorphic computing. Biophysical neuron models are inherently inefficient to emulate on event-driven neuromorphic hardware or robotic platforms. A point neuron model such as the one used in this study requires a minimal set of parameters and only needs to communicate upon firing an action potential, therefore fulfilling the criteria of communication sparseness and memory locality that is a prerequisite for efficient operation on fully distributed neuromorphic computing approaches. Izhikevich models have been successfully employed in various applications on neuromorphic hardware [26, 32, 33] and robots [21, 28, 34]. Temporal dynamics are naturally embedded in the spiking behaviour. The input signals are discretised through spike timing, encoding time-dependent features such as slope. This enables direct integration with event-driven systems without additional temporal encoding or signal transformation. This work supports practical applications such as rapid prototyping, resource-constrained robotics, and educational tools, broadening access to neuromorphic methods without relying on complex biologically detailed systems.

Future research can implement our detectors in various domains where efficient neuronal computations are essential. We will extend our study by creating a neural network of these slope-detecting neurons for gas-based control in mobile robot applications. Efficient models that balance biological realism and information processing is a requirement in dynamic environments. The instantaneous dynamics of gas concentration in a turbulent plume are extremely complex [4, 12], and this rich temporal structure and spatial distribution of gas plumes demand rapid, low-latency responses to temporal cues in these signals [4, 13, 22]. In real-time gas-based control tasks, computational resources, power, and response time are essential. The regular spiking point neuron used in our research functions effectively as a slope-detector, and its simple dynamics make it well-suited for this real-time application. However, we consider extending feature research to alternative models, for example the Adaptive Exponential Integrate-and-Fire (AdEx) model, which also incorporates adaptation [17, 27, 29].

Amplitude and slope are important dynamical features of sensory signals, encoding information about the scene [12, 14, 15]. We plan to develop a network of our slope-detectors to infer direction for odour source localisation. We will implement our bio-inspired solution onto a mobile robot, creating gas-based control in real-time. We believe that using these detectors will advance gas-based control research, by providing a solution to decode key dynamical features of sensory input in a manner suitable for future real-time applications in robotics and neuromorphic hardware.

## Method

All simulations were created using PyNN and conducted using the NEURON simulator [3] with the built-in Izhikevich neuron model [7]. For the extensive search of the Izhikevich parameter space, simulations are performed on the UH-HPC cluster for efficient computational handling. The datasets generated and/or analysed during the study are available in the GitLab repository gitlab.com/r-miko/slope-detector-project, along with the code and a guide on how to run the simulations.

### Identifying slope and amplitude-detectors in the Izhikevich parameter space

We conduct a systematic search of the Izhikevich parameter space to identify sets of parameters that produce either slope or amplitude detecting neurons. The parameters *a, c*, and *d* are investigated over a wide range of values, while *b* is fixed to 0.2 based on previous research [7] and its ability to reproduce desired behaviours. To define the search boundaries, we examine parameters known to produce specific neural dynamics, such as intrinsically bursting or fast-spiking. Simulations are performed on the UH-HPC cluster for efficient computational handling.

Each neuron is injected with a rectified 4 Hz sinusoidal input signal. The spike trains are extracted and compared to the original input signal to determine the features that trigger the neuron’s response. slope-detectors are identified by spike rates corresponding to the rising edges of the input signal, while amplitude-detectors are characterized by spike rates at signal peaks. To quantify detection performance, we calculate the detection percentage, calculated as the fraction of relevant spikes relative to the total spike count, multiplied by 100. For example, slope detection percentage is defined as:

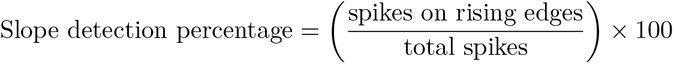

We then examine the relationship between bursting behaviour and detection capabilities. Bursting is defined in this study by an inter-spike interval of 10 ms, and burst percentage represents the proportion of spikes within bursts.

To assess the robustness of the detectors, Gaussian white noise signals are generated, low-pass filtered using a Butterworth filter (5 Hz; *µ* = .006; *σ* = .015), and injected into the detectors. A robust detector should respond consistently across different input types, not just sinusoidal signals.

### Detection frequency range

After creating a slope-detector from the parameter search, we evaluate its response across a range of input frequencies. We generate a Gaussian white noise signal, applying a Butterworth low-pass filter (*µ* = .006; *σ* = .015), and inject it into the slope-detector. This signal is filtered with frequencies between 5 Hz to 30 Hz. We then apply reverse correlation techniques (spike-triggered average) to identify the features of the input signal triggering its response, by aligning the input signal with spike occurrences. By averaging across multiple spikes, it reveals the average input signal associated with the detector’s spiking activity, providing insights into the specific characteristics driving the detector’s response.

### Bi-directional slope detection

We investigate whether the slope-detector is capable of bi-directional slope detection. We observe whether it could detect both the positive slopes of the input signals and the down strokes when inverted. To examine the excitatory response, a Gaussian white noise signal (5 Hz; *µ* = .008; *σ* = .015) is generated and directly inject it into the detector. To simulate the inhibitory input, the sign of the signal is inverted, and the resulting signal is injected into the neuron to observe the inhibitory response.

### Burst duration encoding

We investigate whether the bursts from the bursting slope-detector neuron only marks the occurrence of signal upstrokes or if they also signal the slope magnitude. We analyse the relationship between input slopes and burst durations, where burst duration is defined by the number of spikes in a burst (using a fixed inter-spike interval of 10 ms).

A Kernel Density Estimation (KDE) plot is created to display the number of spikes per burst divided by a Gaussian density estimation, providing the fraction of bursts for different signal slopes. From this, we create a ROC (Receiver Operator Characteristic) curve to observe the discriminability between burst length distributions and use the composite trapezoidal rule to calculate the AUC (Area Under the Curve) value. An AUC of 1 indicates perfect discrimination between the burst length distributions, meaning there is no overlap and the burst length informs the magnitude of the slope.

### Bursting slope-detector comparison

We compare the Izhikevich bursting slope-detector neuron to a two-compartmental biophysical neuron model [5]. The biophysical neuron model consists of a dendrite compartment responsible for bursting behaviour and a somatic region generating fast spikes. To ensure a fair comparison, we create a robust neuron that demonstrates 100 % slope detection percentage and 100 % burst percentage, as identified in the previous analyses. We then ensure that similar input signals are injected by analysing the features and patterns of their inputs and mirroring them. We also create figures in similar formats to allow for a side-by-side comparison.

## Author contributions statement

R.M. Conceptualisation, Methodology, Software, Investigation, Visualization, Writing - Original Draft, Writing - Review and Editing.

M.M.S. Methodology, Writing - Review and Editing, Supervision.

V.S. Conceptualisation, Methodology, Writing - Review and Editing, Supervision.

M.S. Conceptualisation, Methodology, Writing - Review and Editing, Supervision, Funding Acquisition.

## Data availability statement

The datasets generated and/or analysed during the current study are available in the GitLab repository gitlab.com/r-miko/slope-detector-project

